# The brain time estimation system is dynamically assembled by non-temporal perceptual states

**DOI:** 10.1101/2023.09.20.556438

**Authors:** Wenhao Han, Tingting Feng, Yifei Han, Yawen Sun, Yi Jiang, Tao Zhang

## Abstract

To investigate how neural integration of visual signals distorts human time perception, we developed a novel apparent motion paradigm. We found that the higher the frequency of visual events defined by spatial location, the longer the perception duration. At the same visual event frequency, perceptual duration was shorter when motion perception was evoked. We propose that the non-temporal perceptual states dynamically determine which core brain region (read-out system) and which group of activated neurons (timing units) assemble into a time estimation system. Timing units have activation delays in integrating input signals, which results in shorter perceptual time since perceptual duration depends only on the outputs of timing units. The read-out system consumes operating time to accumulate each timing unit, this results in a longer perception time the more timing units are accumulated in the same physical duration. Subsequently, experiments further validated two predictions of our perceptual assembly model.

## Introduction

Timing is critical to all aspects of our lives and brain function, from regulating physiological activity, sensory information processing, decision makings, to motor behavior control. This is why how our brain estimates time is a question that has fascinated researchers in various fields for decades. As an information processing system based on the electrical activity of neurons, how our brain represents time on the millisecond to second scale, termed interval timing, is particularly important for it to perform most functions. Interestingly, although the science and technology created by humans have been able to produce very accurate and stable timekeeping instruments, the human brain’s estimate of time is very imprecise and susceptible to change by various non-time factors compared to clocks, such as many stimulus properties[1], motor behavior[2] and even cross-modality integration[3]. Researchers usually refer to the perceptual time becoming longer relative to the reference duration as a time dilation effect, and vice versa as a compression effect. Over the past century, countless time dilation and compression effects induced by a variety of different non-time factors have been reported. However, the underlying neural mechanisms have never been clearly explained. In a journey to explore the neural basis of time perception, recent advances have proposed that timing is an intrinsic property of most neural circuits. The perceptual time is context/state-dependent and emerges from distributed and inherent dynamics of neural circuits[4–7]. Although the intrinsic model appears to be consistent with the variability of time perception and is neurophysiological plausible in principle, a clear and unified mechanism to demonstrate how exactly various time dilation and compression effects emerge from neural circuits is still lacking.

We attempted to approach such a goal using visual motion as the probe, where the time variable is essential for both neural computation and perception. The brain network model studied was also limited to feedforward processes in a series of brain regions from the retina to the dorsal pathway of the visual system. The visual stimulus is a point of light that is presented over time at different spatial locations in an invisible circle around fixation point (Fig. 1a). It would induce an apparent motion perception when presented in successive spatial locations over time. It is worth pointing out that in the laboratory environment, the visual display presents all stimuli as a series of static images with different refresh rates, rather than continuous motion as in the real world. Previous evidence also suggests that apparent motion can effectively activate both V1 and MT and evoke visual motion perception [8]. The apparent motion has a clear advantage regarding event control of sensory information input because, from the perspective of retina photoreceptors, each flash (dot presented per frame) can be considered as an independent and position invariant visual event. This allows us to tightly control when and which spatial location information is received by the visual system at the source of information input, and thus better examine how temporal integration of spatial locations affects our time perception. Our hypothesis is that in the absence of evoked visual motion perception, perceptual time depends primarily on the activated spatial location-sensitive neurons, whereas in the presence of evoked visual motion perception, perceptual time depends primarily on the activity of those neurons that temporally integrate the output of the spatial location-sensitive neurons.

**Figure 1.**
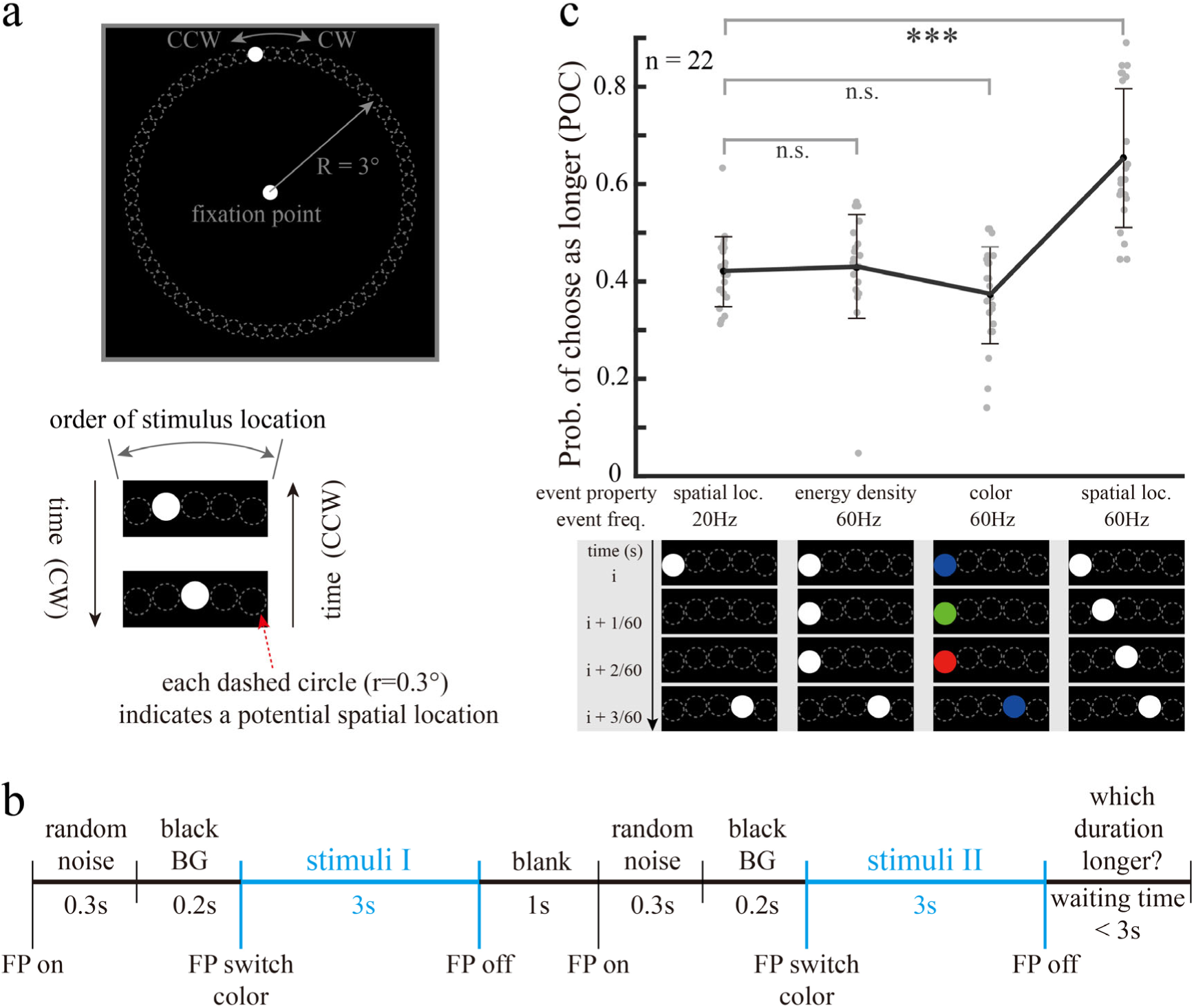
**a.** The visual stimuli were successively presented white dot along a full circle path centered at the fixation point. The moving direction (the apparent rotation direction) was counterbalanced. **b.** The diagram of the experimental procedure. Participants pressed the answer key at the end of each trial to report which of the two successively presented stimuli with the same duration lasted longer perceptually. FP, fixation point; BG, background. The random noise and black BG right before each stimulus were used to set the active state of the neural network to the same level. For each experiment, 25% catch trials with real time differences (the presentation durations of the two stimuli were significantly different in each trial) were included to assess the subject’s effort in doing the experiment. **c.** In each trial, two of the four visual stimuli with different event properties (labeled below the x-axis) were randomly selected. The subjects were asked to judge which stimulus evoked the longer time perception. The temporal sequence of flash dot positions was illustrated at the bottom. POC: mean_spatial loc. 20Hz_±SD=0.45±0.08, mean_energy density 60Hz_±SD=0.46±0.11, mean_color 60Hz_±SD=0.40±0.11, mean _spatial loc. 60Hz_±SD=0.70±0.15.

We adopted the classical duration comparison experimental task, where subjects had to report which of the two successively presented stimuli lasted longer at the end of each trial (Fig. 1b). Before each stimulus presentation, we added a white noise stimulus followed by black background for the purpose of eliminating visual after-effects and setting the visual system in identical network activity status.

## Results

First, we want to find out which event property is crucial for time perception. The event properties tested were selected based on the coding ability of primate retinal neurons (Fig. 1c). Comparing to the reference condition (20Hz), the same retina ganglion cells in the same time interval received triple light quanta under energy density condition (60Hz). Similarly, different color sensitive photoreceptors at the same retina location were activated sequentially under color condition, but the overall energy is roughly the same as reference condition. However, under spatial location condition, triple amount retinal ganglion cells were activated successively because of the two additional stimulus locations. We use probability of choice (POC) to quantify the relative perceptual duration, which is the proportion of choose one stimulus condition as longer among all trials that included this stimulus condition (see Methods for details). Therefore, the larger the POC, the longer the perception time. We find that increasing the number of spatial locations leads to time dilation (Fig. 1c, t-test, t = 6.23, p < 0.001), while increasing the number of presentations and color changes has no effect on perceptual time interval. The results suggest that spatial location is the key event property affecting time perception for time perception evoked by visual signals. This is consistent with previous findings that time distortion is spatially localized [9], spatially selective [10] and spatially specific [11]. As the basic information we acquire directly from the external world, spatial location should be the most fundamental building block for the neural representation of visual motion.

Before we systematically assess the time distortions caused by spatiotemporal integration, a baseline for perceived time should be established. For this purpose, there is no better baseline than temporal perception induced by spatial location cue alone. Therefore, we randomized the order of spatial locations of flash dot so that the change in time perception depends only on the number of spatial locations. In contrast to the spatial location condition in previous experiment, this manipulation excluded the involvement of spatial-temporal integration neural circuits (Fig. 2a, bottom), at least those neurons that integrate local spatial locations. The event frequency here is defined as the number of different spatial locations presented per second. We find that perceptual time becomes longer with increasing event frequency, implying that increasing the number of spatial locations can induce time dilation (Fig. 2a, top, t-test, 30Hz vs 60Hz, t = 3.09, *p* = 0.013). Because larger stimuli activates more spatial location sensitive neurons, this result is actually consistent with previously reported time dilation effect cause by increasing stimulus size[12].

**Figure 2.**
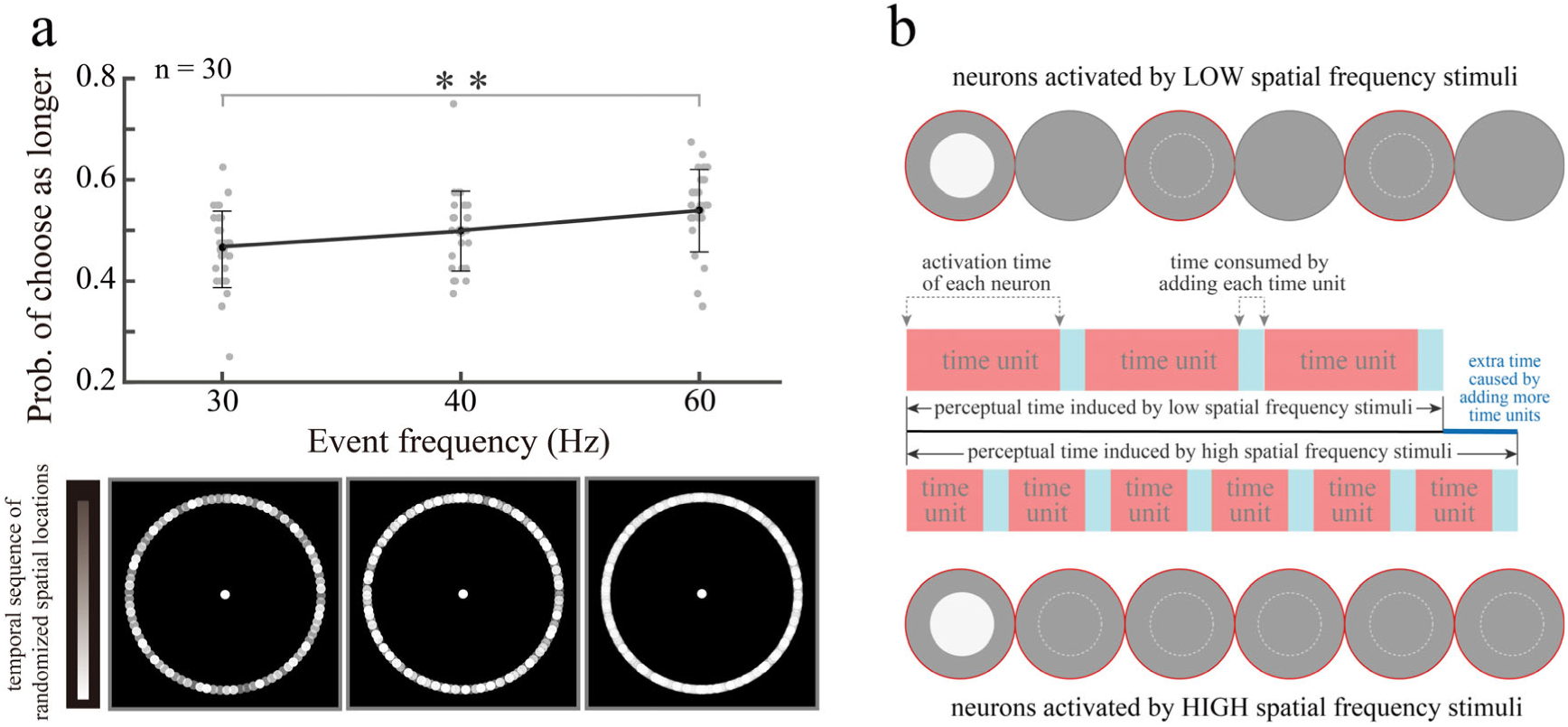
**a.** The visual stimuli were single flash dot presented at randomized spatial locations. The Y axis denotes the probability of choosing this event frequency stimuli as longer. POC: mean_30Hz_±SD=0.46 ±0.08, mean_40Hz_±SD=0.50±0.08, mean_60Hz_±SD=0.54±0.08. **b.** The gray discs indicate the receptive fields of the neurons, the white dots on them indicate the visual stimuli, and the light gray dashed circles indicate the possible spatial locations of the visual stimuli. The gray discs with red edges indicate neurons were activated during stimulus presentation. For the case illustrated here, the high spatial frequency visual stimulus activated twice as many neurons as the low spatial frequency condition, but the activation time of the neurons corresponding to each spatial location is only half of the latter (each dot presented 2 frames under 60Hz condition, but 4 frames under 30Hz condition, see Methods for details).

We propose a possible explanation for the underlying neural mechanism of this result (Fig. 2b). If we define the activation of neurons at the same spatial location as a timing unit (the basic unit for measuring temporal duration), then the perceptual duration can be considered as the sum of all timing units activated by visual stimuli, just like the working principle of a clock. The summation operation is performed by neurons in a core brain region (termed read-out system/neurons) that receives all timing unit projections. However, what is often overlooked is that each summation operation consumes a fraction of time! The same principle applies to both man-made clock and “timing device” in our brain. For the same stimulus duration, the higher the event frequency, the more spatial location detection neurons are activated by the visual stimulus (the shorter the activation time of neurons at each spatial location), and the greater the number of summation operations, the more time they consume. Because perceptual time is composed of both timing units and the consumed time by read-out system to sum them, this leads to the phenomenon that high event frequency stimuli induce longer perceptual time. This effect should have an upper bound. When two successively presented visual stimuli are so close in spatial location that the neurons cannot distinguish between them, increasing the event frequency of the stimuli will no longer result in longer perceptual times.

We then examined the influence of spatiotemporal integration neural circuits on perceptual time, by pairwise comparing the effects of positionally random and successively presented stimuli at the same event frequency (Fig. 3a). Relative to the random spatial location condition, perceptual time becomes shorter when flash dot successively presented, and the effect appears to be quite stable across different event frequency (POC[30Hz, 40Hz, 60Hz] vs 0.5, one sample t-test, t = 3.17, p = 0.002). This result suggests that perceptual time becomes less after the involvement of spatiotemporal integration neural circuits. It seems some time has been “stolen” by the temporal integration of spatial locations!

**Figure 3.**
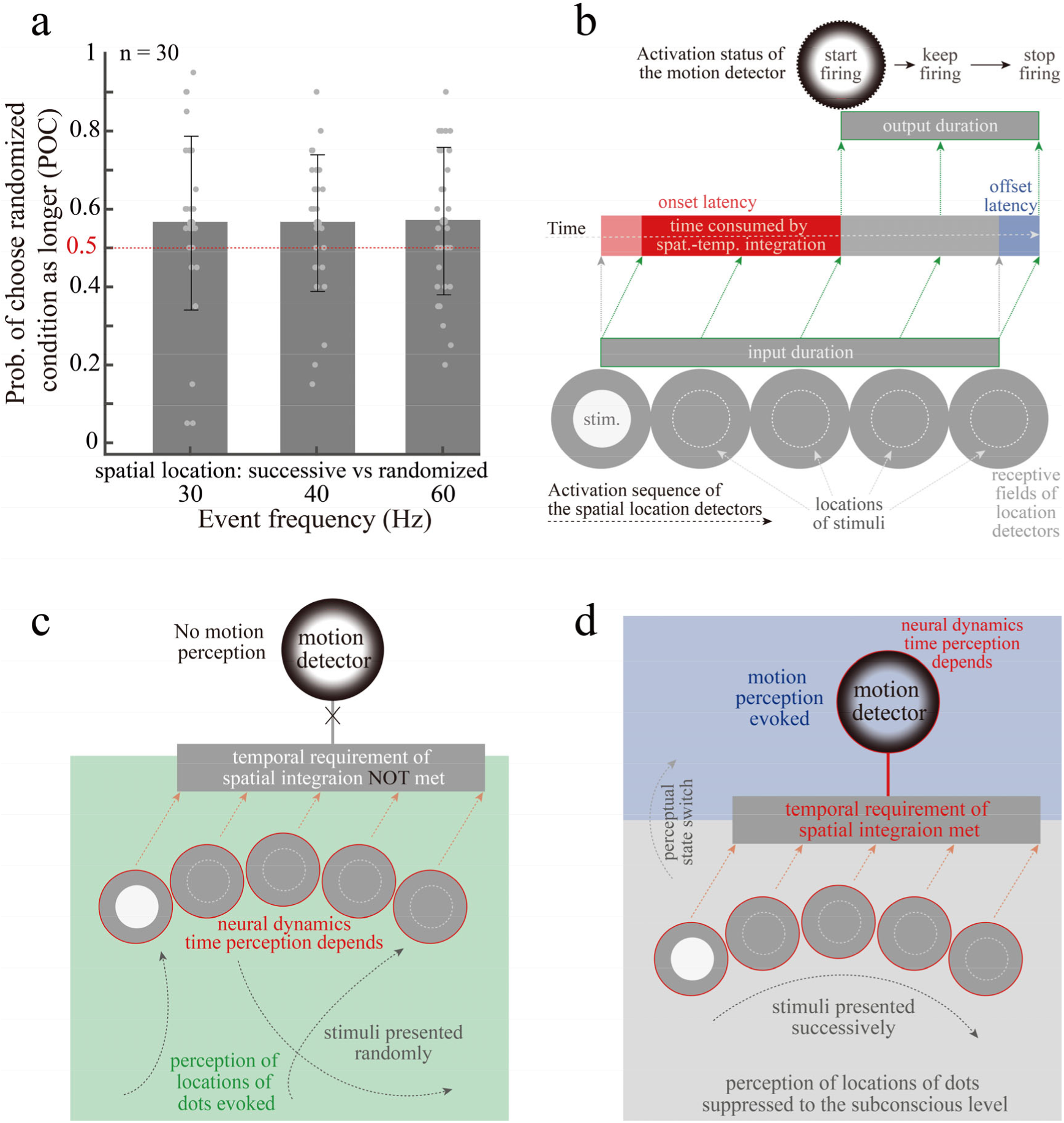
**a.** The two visual stimuli included in each trial remained identical in both spatial locations and the presenting frequency (same event frequency). The only difference was that the presenting order of white dots was randomized for one and spatially continuous for the other. POC: mean_30Hz_±SD=0.56± 0.22, mean_40Hz_±SD=0.56±0.18, mean_60Hz_±SD=0.57±0.19. **b.** Illustration of how time is stolen by temporal integration of spatial locations. The input duration is the overall activation duration of all spatial location detection neurons that project to the same motion detection neuron. The onset latency contains two components, a transmission delay (pink) of the signals from the spatial location detection neurons to the motion detection neuron, and a signal integration delay (red) required to accumulate sufficient input signals to activate the motion detection neuron. The output duration is the time that the motion detection neuron is activated. **c,d.** Illustration of the perceptual state dependence of time perception.

To reveal how perceived time is shortened by spatiotemporal integration, we analyzed the process of temporal integration of spatial location signals by motion-sensitive neurons (Fig. 3b). As we know, each motion-detecting neuron receives projections from multiple location-detecting neurons. When a visual stimulus activates these location detection neurons sequentially and in temporal order along the preferred direction of the motion detection neuron, the spatial location signals arrive the dendrites of the motion detection neuron in the same temporal order. For motion detection neuron, EPSPs (excitatory postsynaptic potentials) generated at synapses farther from the cytosol take longer to transmit to the cytosol, membrane potentials are superimposed when multiple EPSPs arrive at the cytosol at the same time, and if the superimposed membrane potentials exceed the excitability threshold of the neuron, the motion-detecting neuron is activated[13]. Therefore, the dendrites of a motion-detecting neuron can be considered as a temporal structure that integrates the input signals[14], and a direct evidence is that single cortical pyramidal cell dendrites can encode temporal sequences of synaptic inputs[15]. Note that in order to reach excitability thresholds, motion-detecting neurons need to integrate multiple temporally continuous spatial location signals for the obvious reason that motion signals cannot be generated by a single spatial location signal. This part of the integration time consumed to bring the motion-detecting neuron to the excitation threshold (Fig. 3b, red bar) is not reflected in its output duration. This results in a shorter duration of excitatory output from the motion-detecting neuron than the duration of the spatial location input signal it receives, which in turn is approximately the same as the duration of visual stimulus with a slight offset delay caused signal transmission. The neural mechanism we proposed here is supported by previous findings that the onset latency is generally longer than offset latency in brain regions that process visual motion[16]. Thus, the perceptual time became shorter when we used the output duration of motion-detecting neurons as a timing unit.

One may ask why our brain does not use the activation time of spatial location detection neurons as timing units, they are also activated under successive stimulus condition. We did use the activation duration of the spatial location neuron as the timing unit in the random stimulus condition (Fig. 3c), but when motion perception was evoked, the timing unit switched to the output duration of the motion detection neuron (Fig. 3d). In the random stimulus condition, the only timing units available were the output durations of the location detection neurons. If the same timing units were still used in the successive stimulus condition, the perceived times should be the same because the event frequencies are the same. However, it turned out that the perceived time was significantly shorter after motion perception was evoked, indicating that perceptual state has an important effect on perceptual timing. We hypothesize that the timing unit used for time estimation is the neuronal excitation that directly supports the current perceptual state. That is, the current perceptual state determines which set of activated neurons is used as the timing unit.

Our previous result demonstrates that increasing event frequency can lead to longer perceptual time (Fig. 2), but this result is in the case where motion perception has not been evoked and the timing unit is the activation state of the spatial location detection neuron. Does this rule remain valid when motion perception is evoked, and the timing unit becomes the activation state of the motion detection neuron? We thus measured the dependence of time perception on the frequency of spatial location events, when the white dot stimuli were presented successively in term of both temporal sequence and spatial order. Because subjects no longer perceive independent events (i.e., flicker of light dots) when the event frequency increases to a certain point, we also defined a continuous motion sensory threshold to describe the minimum event frequency required for subjects to perceive continuous motion without flicker and measured the threshold for each subject using an independent procedure prior to the formal experiment (see Methods for details). We first examined the dependence of perceived time over a wide range of event frequencies (Fig. 4a, from 15Hz to 120Hz at double rate). We find that when not exceeding 60 Hz, perceptual time increases continuously with spatial frequency (Fig. 4a, multiple t-test, 15Hz vs 30Hz, t = 9.26, *p* < 0.001; 30Hz vs 60Hz, t = 4.73, *p* < 0.001), which means the number effect rule (more timing units, longer duration perceived) still hold in the state of motion perception.

**Figure 4.**
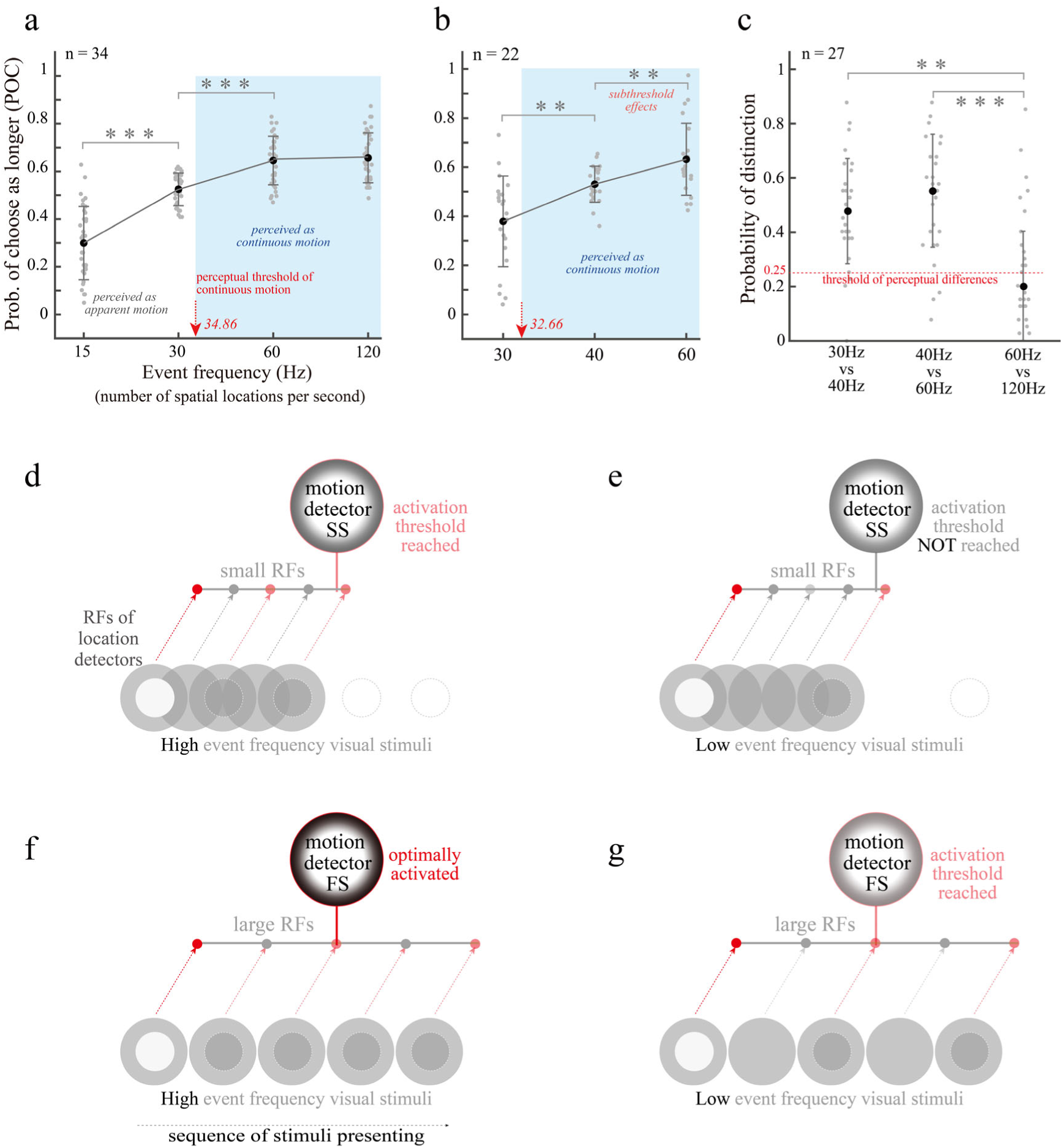
**a,b.** How perceptual time is affected by event frequency. The red arrows indicate the perceptual threshold from apparent motion to continuous motion. **c.** Subjects were asked to report by keystroke whether there was a perceptually difference between the two visual stimuli in each trial. The probability of distinction is the percentage of trials in which the subject can perceive a difference over all trials in that comparison condition (see Method for details). The black dots represent median instead of mean. The threshold of perceptual differences is an arbitrary value representing 25% probability that the subject will perceive the stimulus difference. **d∼g.** Illustration of how different event frequencies affect the activation of different types of motion detectors.

As you can see in Fig. 4a (red arrow), the perceptual threshold of continuous motion is slightly less than 35Hz. Because subjects are subjectively unable to distinguish between independent photopic stimulus events when the presentation frequency of white dots exceeds this threshold, this threshold can be considered as the dividing of perceptual awareness line between apparent motion and continuous motion. It is interesting to know whether the number effect rule still exists above this threshold. It does (Fig. 4b, multiple t-test, 30Hz vs 40Hz, t = 2.95, *p* = 0.008; 40Hz vs 60Hz, t = 3.40, *p* = 0.005)! This result suggests that, at least for the sensory time estimation induced by visual signals, there is not much difference between apparent motion or continuous motion as long as visual motion perception can be induced. From another perspective, perceptual timing relies on the perceptual state rather than the stimulus form.

Another noteworthy aspect is that the perceptual time no longer increases when the event frequency exceeds 60 Hz (Fig. 4a), indicating the very existence of a maximum effect limit as we expected before (the upper boundary). We speculate that this maximum effect limit is consistent with the discriminative limit of motion perception. We tested this hypothesis in a new group of subjects. The visual stimuli were the same as that used in Figures 4a and 4b, but the experimental task was for subjects to determine whether there was a perceptual difference between the two stimuli in each trial (not a temporal difference!). We found that subjects were perceptually unable to distinguish between the 60Hz and 120Hz conditions, but could perceive differences between the 30Hz vs 40Hz, and the 40Hz vs 60Hz conditions (Fig. 4c). This result coincides with the changes of time perception evoked by visual motion (Fig. 4a). It suggests that when the event frequency increases to the point that it no longer induces perceptual differences, it will no longer affect our time perception either.

Did high event frequency visual stimulation here indeed activate more motion detection neurons? It should be the case. The visual stimuli with different event frequencies characterize the same spatio-temporal relationships in terms of moving direction and speed. The difference is that visual stimuli with low event frequencies provide sparser spatial location signals. We know that even neurons that perform the same function in the same brain region have different receptive field sizes. For example, the size of the receptive fields of those neurons encoding the direction of movement in the MT brain region of macaques can vary several times. Previous studies have found that the spatiotemporal integration patterns of speed-tuned neurons in MT are diverse [17], meaning that even if a group of neurons has the same speed selectivity, their preferred spatial frequencies will be different. Under high-event-frequency stimuli (Fig. d&f), motion-detecting neurons with both large and small RFs can be activated. Under low-event-frequency stimuli (Fig. e&g), some neurons with small RFs cannot be activated because of insufficient spatial location signals received (Fig. 4e), which means that fewer motion detection neurons are involved in time estimation.

To summarize our findings, we illustrate them in a unified framework, which we call the perceptual assembly model (Fig. 5). It can be expressed in detail as follows:

1. Perceptual state dependence principle (Fig. 5a). Time estimation is only a function of perceptual neural networks. The time estimation system of our brain is composed of a group of timing units and a read-out core brain area which connects with them. Those two components are dynamically assembled based on the perceptual state. The timing units refer to activated neurons directly supporting the current non-temporal perceptual state. Please note that the perceptual intensity/strength can vary within the same perceptual state. For example, MT neurons with same direction selectivity and same speed preference corresponds to same perceptual state, but how many of those neurons are activated determines perceptual intensity/strength.
2. Time decremental law of signal integration (Fig. 5b). The perceptual state is dependent on the output of the corresponding activated neurons. Because the temporal variation due to the signal integration process below current perceptual state is not accounted for in the time estimation, this results in the signal integration time consumed to reach the activation threshold being excluded from the perception time, which in turn leads to a relative time compression effect. The more layers of signal integration, the longer it takes to reach the activation threshold (minimum onset latency of brain regions involved in this study: LGN 20 ms [18]; LGN magno. 16 ms [19]; V1 28 ms and MT 39 ms [20]; V1 34 ms, MT 49 ms and MST 58 ms [21]), and the shorter the corresponding perception time. For a given modality, the longest perceptual time is determined by the frequency of the largest event that can induce a difference in perception without temporal integration of the original input signal.
3. Time incremental law of perceptual summation (Fig. 5c). The read-out system consumes a little bit of time for each summing operation of the timing unit. Because this summation is in the perceptual state, it is included in the perceptual time. This results in longer perceptual time as the number of summed timing units increases. This effect has an upper limit, i.e., the activation of additional timing units will not be included in the summation calculation if they can no longer cause perceptual changes.

**Figure 5.**
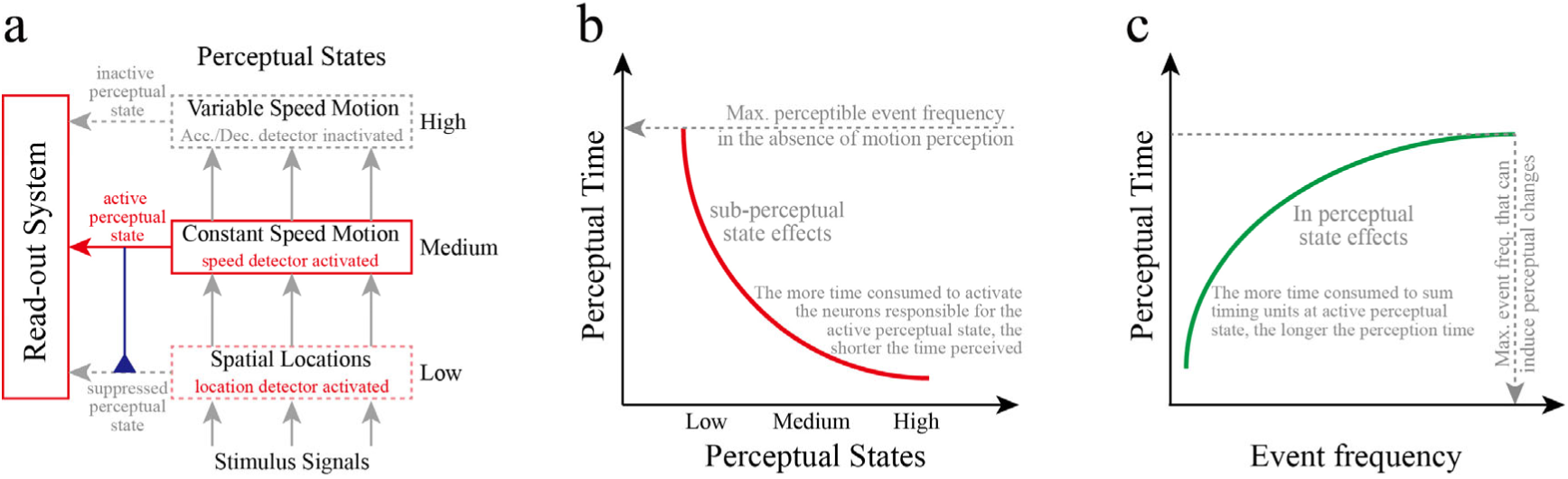
**a.** Perceptual state dependence principle. This principle in fact describes the dynamic mechanism of the composition of the brain timing system. The active perceptual state determines which population of activated neurons constitutes the “timing device” with the read-out system. The high, medium and low on the right side refer to the information processing hierarchy. For clarity, here we only graphically illustrate how the perceptual state dynamically assembles the time estimation system between different hierarchy levels. **b**. Time decremental law of signal integration. It describes the law that, for the same stimulus duration, the perceptual time becomes relatively shorter as the level of signal integration corresponding to the current perceptual state increases. For a specific type of signal input, such as the spatial location signal here, the maximum perception time is determined by the highest perceptible event frequency without temporal integration of input signals (no visual motion perception evoked yet). We believe that the relationship between perceptual time and perceptual state should be nonlinear as shown in the figure. There are two reasons: first, the integration of dendrites to input signals is nonlinear; second, the cross-layer projection can narrow the onset delay of higher order brain areas relative to lower ones to some extent. **c**. Time incremental law of perceptual summation. It describes how the perception time becomes relatively longer if the read-out system needs to accumulate more timing units for the same stimulus duration at same perceptual state. For a specific perceptual state and stimulus duration, the longest perception time is determined by the maximum event frequency that can induce perceptual changes. We express the relationship between perceptual time and event frequency as a nonlinearity as shown in the figure. This is only one possibility. Another possibility is that the relationship may be linear after the event frequency is high enough to elicit stable perception, but before the highest perceptual discrimination accuracy has been exceeded.

Notably, both laws describe how perceptual time varies under the same stimulus duration conditions, which means that our model focuses on describing the neural mechanisms of perceptual time distortion and the corresponding laws, rather than describing how perceptual time varies with physical time, as most other models do. Although this model was constructed based on the neuronal processing of motion information from visual signals, we believe it can be applied to other modalities, such as auditory and tactile. It is important to note that, tactile signals are spatially localized but auditory signals are not, probably that’s why the time distortion effect induced by sound cues relies more on stimulus intensity instead of temporal integration of spatial locations [22]. This framework is based on the feedforward visual motion information processing in hundreds of milliseconds to seconds range, it might be invalid if long-distance feedback regulation, attention modulation and memory were involved, because they can either interfere with the dynamic assembly relationship or change the number and output of timing units.

A straightforward prediction of our model, for the influence of timing units and perceptual state dependence, is that the time-compression effect caused by spatiotemporal integration accumulates as the level of integration increases, if higher levels of integration depend on the direct output of lower levels. Neurophysiological evidence from the macaque visual system supports this hypothesis, as the response latency of neurons does increase progressively with the level of signal processing[21]. This prediction can be examined by comparing the perceived duration under acceleration and deceleration with the perceived duration in the constant speed condition, since the former can be seen as the temporal integration of the speed signals.

Let’s consider how neurons calculate speed and how speed changes affect perceptual timing first. As we know, some brain regions (for example, MT [23]) that process visual motion have neurons that prefer different speeds. A convenient way to generate speed selectivity is to take advantage of the flexibility of the temporal structure of neuronal dendrites, that is, the difference in dendrite size. Suppose we have a neuron with a preference for moderate speed that integrates spatial location signals as shown in Fig 6a (top). By aligning the synaptic sites on the dendrites that receive the same spatial location signal more closely, we can make this neuron more likely to be activated by the same displacement produced in a shorter period, i.e., a preference for faster speed (Fig. 6a, middle). A direct consequence of this is that the time consumed to integrate the same spatial locations also becomes shorter. The construction of neuron that prefer slow motion is similar, except that the time consumed to integrate the information becomes longer (Fig. 6a, bottom). The direct evidence for this hypothesis is that neurons tuned to slow speed has longer latency, which means that more time is needed to integrate same amount of spatial locations to reach activation threshold, than those tuned to fast speed [24] and the speed detection ability of neurons indeed critically depends on their dendritic geometry [25]. According to our model, because neurons that prefer fast motion consume less time in integrating spatial location signals than neurons that prefer slow motion, fast motion stimuli will induce longer perception time than slow motion stimuli. This is exactly the case [26]!

**Figure 6.**
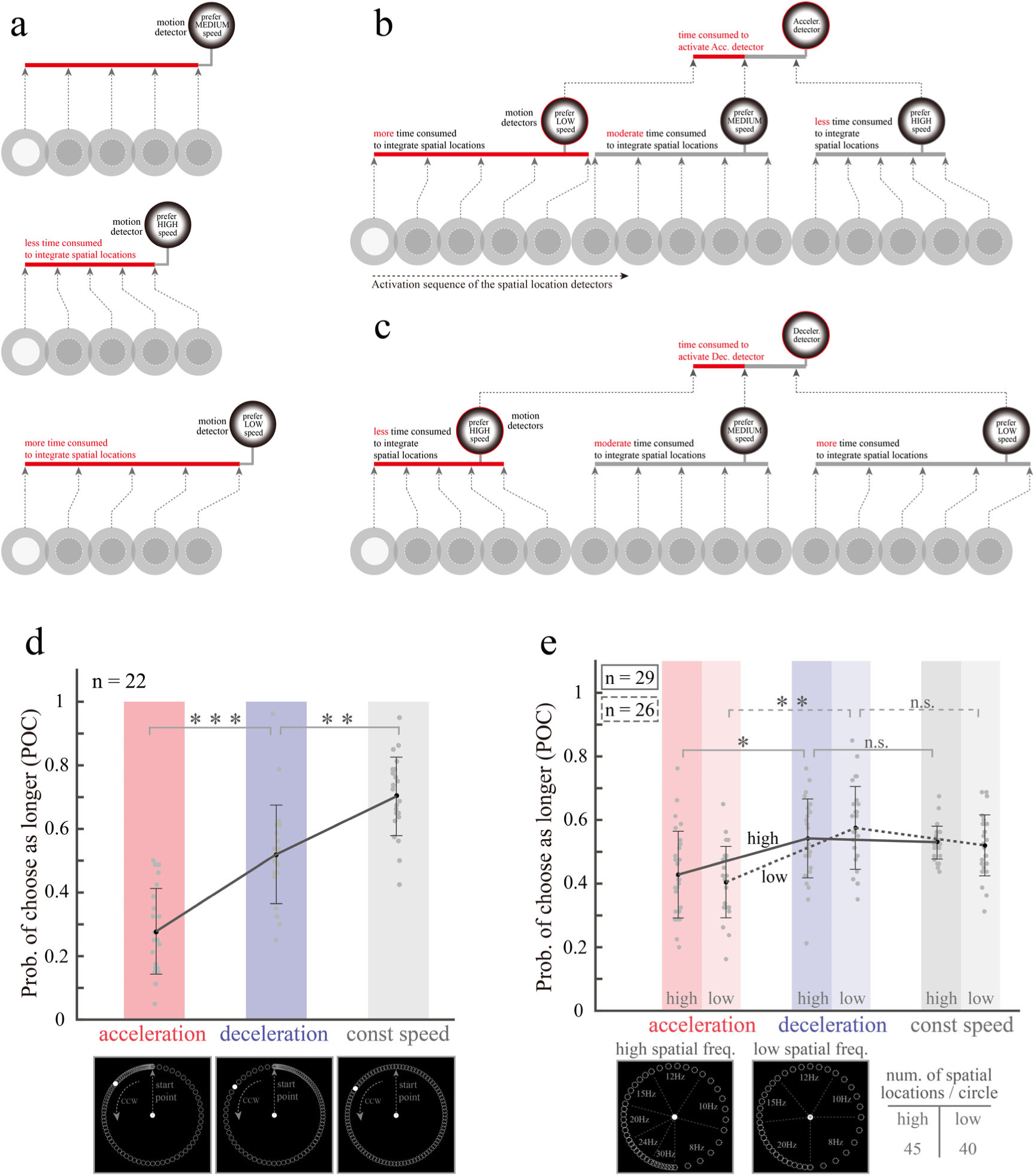
**a.** How to build motion-detecting neurons with different speed preferences by changing the integration time. **b,c.** The proposed dendritic computational structures for acceleration and deceleration detectors. Both acceleration and deceleration detectors calculate the rate of change of the velocity signal over time, the only difference being the opposite trend of change. Here we assume that they consume the same amount of time to reach activation threshold in the stage of integrating speed signals (top red bars in Fig. b&c). **d.** How perceptual timing varies with motion perception states. mean_acceleration_ ±SD=0.278±0.135; mean_deceleration_±SD=0.520±0.155; mean_const speed_±SD=0.702±0.124. **e.** Perception time can be varied by adjusting the number of activated neurons. High spatial freq. experiment: mean_acceleration_±SD=0.419±0.145; mean_deceleration_±SD=0.552±0.132; mean_const speed_± SD=0.529±0.051. Low spatial freq. experiment: mean_acceleration_±SD=0.405±0.112; mean_deceleration_± SD=0.575±0.130; mean_const speed_±SD=0.520±0.096.

How to construct a neuron detecting speed changes? This can also be achieved by the different spatio-temporal structure of the dendrites receiving information. The acceleration detector first receives slow motion signals from the distal dendrites and finally receives fast motion signals from the synapse near the cytosol (Fig. 6b), the deceleration detector works the opposite way (Fig. 6c). Because the acceleration detector inherits the onset latency of the slow speed detector, which is larger than the onset latency of the fast speed detector inherited by the deceleration detector, acceleration should induce shorter perceptual time than deceleration according to our model. Due to additional time consumed by integrating speed signals, the perceptual time induced by either acceleration or deceleration should be shorter than constant speed (Fig. 6 b&c).

We conducted an experiment to test these predictions. The experimental procedure was same as before (Fig. 1b), but the visual stimuli were acceleration, deceleration, and constant speed apparent motion (Fig. 6d, bottom). We find that the perceptual time appears to be the longest for constant speed and shortest for acceleration (Fig. 6d, top, multiple t-test, const speed vs deceleration, t = 3.48, *p* < 0.01; deceleration vs acceleration, t = 4.31, *p* < 0.001). This result is consistent with previous reports, although the visual stimuli and experimental paradigm used were slightly different [8].

Another prediction of our model, derived from the law of sum of timing units, is that the perceptual time in previous experiment can be shortened by reducing the number of activated neurons. For obvious reason, it is very important to strictly control the number of activated neurons in this experiment. This can be achieved by ensuring that each white dot presented is perceived as independent event. In fact, in the previous experiments, because the speed changes were continuous, the spatial location distribution of the visual stimuli at the end of the acceleration stimulus phase and the beginning of the deceleration stimulus phase was very dense, resulting in some white dots not being perceived as independent events because they were spatially too close to the previous one. Therefore, we varied the apparent motion speed by group of spatial locations and kept the highest event frequency below the sensory threshold of continuous motion (Fig. 6e, bottom). We set the minimum event frequency to 8Hz because previous studies have indicated that for effective activation of motor-sensitive neurons in V1 and MT, the frequency cannot be lower than 8Hz[27]. The pilot test showed that this visual stimulus was effective in inducing motion perception of constant speed, acceleration, and deceleration among human subjects. Theoretically, reducing event frequency has less impact on acc./dec. detectors, because of their significantly larger receptive fields than speed detectors [28]. Two experiments were conducted, one setting the maximum event frequency to 30Hz (Fig. 6e, high spatial freq.) and the other to 20Hz (Fig. 6e, low spatial freq.). Just as we expected, we find that the difference of perceptual time between constant speed and deceleration was vanished (multiple t-test, High spatial freq.: acceleration vs deceleration, t = 2.64, *p* = 0.027; deceleration vs const speed, t = 0.89, *p* = 0.379. Low spatial freq.: acceleration vs deceleration, t = 3.89, *p* = 0.002; deceleration vs const speed, t = 1.40, *p* = 0.173), and even the perceived duration under constant speed condition is slightly shorter than that of deceleration condition, although not to a statistically significant level.

## Discussion

For the past two decades, researchers have begun to focus more on the effects of timing unit diversity [4, 6, 7, 29, 30] and state dependencies [31, 32] on perceptual time variability. However, a clear and neurophysiological plausible explanation is still lacking to systematically elucidate how exactly the neural dynamics affects perceptual time. It is in this regard that the present study seeks to contribute. We have constructed a novel experimental paradigm for the study of time perception, whose greatest advantage is that it allows strict control over the raw signal input to the brain neural network and its time-series characteristics. Because the concept of motion itself is an ordered temporal integration of spatial signals, and we have a relatively clear understanding of the brain information processing mechanisms of visual motion, it is an ideal working model for studying the neural mechanisms of time perception. We first identified spatial location as the most critical raw input signal for time perception, and then found that the neuronal integration of spatial location shortens perceptual duration, while increasing the number of activated neurons directly related to the perceptual state increases perceptual duration. Hereby we propose a neurophysiological plausible perceptual assembly model to explain various perceptual time distortions in a unified framework (Fig. 5). The model is conceptually simple and somewhat analogous to clock timing, consisting of timing units, accumulators (readout systems, the hippocampal [33–35] for example), and a perceptual dependency that flexibly connects the two. Merchant et al. [36] once pointed out that a core timing mechanism interacting with context-dependent areas maybe the neural basis of the perception and estimation of time. Our findings seem to fit well with this insight.

### The perceptual state dependence principle

Although perceptually we are closely integrated with the external world, the brain is well isolated from the external world and even from our internal environment by layers of bones and tissues. Perceptual experience is nothing more than an internal neural representation of the external world constructed by the brain’s neural networks through the processing of signal input from various receptors. Interestingly, although time is critical for performing normal brain functions and controlling behavior, we do not have time receptors, meaning that subjective time estimates must rely on the structural basis or operating principles of the neural networks. In fact, time is never an independent concept, nor does it have an absolute meaning. It arises from our need to describe the sequence of events, the continuity of events, and the interval between events. For this purpose, humans create timing devices by accumulating events with relatively fixed duration. So essentially measuring time with a clock/timer is using the number of stable events in a series to quantify other events and their intervals of occurrence. We believe that the brain estimates time in a similar way. The pacemaker-accumulator models were developed based on the same idea[37, 38]. However, unlike a clock using a single accumulator to summarize the number of events generated by a single oscillator, we propose that the brain flexibly assemble an accumulator and a group of timing units into a task-oriented temporary time estimating system based on perceptual state. This flexible assembly mechanism should be the main reason why our time perception seems to be unstable and less accurate than man-made clocks.

Buonomano and Maass argued that spatial and temporal processing are inextricably linked [32]. Hass and Durstewitz proposed that the representation of interval time may be an inherent property or “byproduct” of other computations performed by various brain areas[39]. We strongly agree with those insights. What we call a time estimation system assembled based on perceptual state is in fact a functional description of existing non-temporal information processing neural network, which is the visual motion processing network in this study. Thus, the capability of time estimation is limited by the non-temporal information processing network too. The results we demonstrated in Fig. 4a and 4c is a good example of this point. When changes in visual stimuli no longer induce changes in motion perception, the corresponding time perception also ceases to change. The same logic actually applies to the world outside the brain. When the physical world ceases to change, does time still have meaning?

The perceptual state dependence we proposed sounds like the well-known state-dependent model of time perception. It should be clarified, however, that perceptual state dependence here only refers to the principle of flexible assembly of read-out system and timing units, and the state-dependent in their model refers to that temporal information is encoded in state-space trajectories derived from the time-varying changes of activated neural network and synaptic plasticity[32]. Hass and Durstewitz [39, 40] proposed several experimental constraints on time perception models. We believe that the pattern of change of temporal perception with respect to stimuli within a given perceptual state should satisfy these experimental constraints.

### Time decremental law of signal integration

The concept of timing unit is not new. Timing units are used in the working principle of clocks and in many models of time perception. David Eagleman also suggests that in studying the neural basis of differences in perceptual timing, we should give more attention to differences in neuronal response latencies[41]. However, a long-ignored fact is that when the activation state of a neuron is used as a timing unit, the integration of the input signal leads to a shorter perception time (Fig. 3a), because we only utilize its output signals to estimate time and the neuron consumes time to reach the activation threshold (Fig. 3b). Our proposed time decremental law for signal integration (Fig. 5b) points to this mechanism and gives its effect on time perception as the level of signal integration increases, as well as a boundary indicator for this effect. We hope this will provide a unified framework for the effects of various time units on time perception.

The dendritic structure of neurons is very important to its’ function, which is relatively stable in a developed sensory system. This is why relatively stable modulated responses can be obtained when repeated measurements are made on neurons using the same stimulus. However, when other modulatory signals intervene, such as attention or emotion, or when dendritic structure changes due to synaptic plasticity, the output signal of the neuron changes, which in turn affects subjective time perception. Buonomano and Maass has basically discussed all the possible influence factors regarding this issue [32].

There are at least two aspects about timing units that deserve further study. One is whether all neurons are eligible to be used as timing units? Since there are hundreds of different functional types of neurons in the brain, we hypothesize that those sensory/perceptual feature detection and motor planning/control neurons that are sensitive to time are more suitable as timing units. Another aspect is which property of timing unit (the output signals of activated neurons) we are using to estimate the time.

We know that the neuron encodes the stimulus signal characteristics in the sequence of action potentials. It has several properties, such as onset moment, activation duration, average intensity (firing rate), etc. So which property of the action potential sequence do we use for timing? The intuitive feeling should be the activation duration, but that may not be the case. This is because the clock integrates time units from a single source, whereas the read-out system of brain integrates time units from multiple sources in a distributed manner. The former ensures that there is no temporal overlap between the successive unit, while the latter is not guaranteed because the previous neuron may not cease firing when the next neuron is activated. From this perspective, the onset moment of neuronal activation should be a better candidate, which we plan to test in the future.

### Time incremental law of perceptual summation

The concepts of accumulator and read-out neuron/system have also been around for a long time. A linear variation of its output signal intensity with time should be its necessary feature, e.g. ramping activity. However, it is not clear how the read-out system interacts with the timing units and further affects time perception. We propose that the read-out system sum each timing unit requires an operation time, which is incorporated in the time perception. This results in that more timing units are summed within a fixed objective time interval, the longer the subjective perception time is, and give a boundary indicator to describe the maximum perception time under this condition (Fig. 5c). It is important to note that this effect is only valid when temporal comparisons are made in the same perceptual state. We hope this will provide a unified explanation of how time perception is affected while varying number of time units. For example, increasing the size of visual stimuli results in longer perceptual time[12]. Although our conclusions are based on the study of visual motion, the above law should hold true for other modalities as well. In that case, denser sound cues, more complex odor or tactile stimulus patterns should all lead to longer perceptual times when testing only in the respective modalities and keeping the perceptual state unchanged.

Although we cannot yet determine which brain regions are read-out systems, it should have the following characteristics. The first is that the neurons or local neural circuits in this brain region should have the ability to integrate input signals linearly, i.e., have time-varying response properties, as mentioned earlier. The second is that this brain region should receive common projections from multiple sensory/perceptual or motor planning/control regions to provide a uniform temporal reference in situations of cross-level perception in same modality, multisensory integration, or sensorimotor synergy etc. The third is that this brain region either has a memory function itself or is closely linked to a brain region with a memory function so that it can temporarily store time interval (or event sequence) for relative temporal comparison. Both hippocampus[35] and cerebellum are good candidates according to these criteria. The cortico-thalamic-basal ganglia is another one, as Merchant et al. suggested[36].

### Sensory signal induced time perception

In studying the phenomenon of time distortion induced by sensory signals, special attention should be paid to the population of neurons activated and the perceptual state evoked, since time estimation is strongly dependent on both. Attentional modulation, sensory modality differences, memory effects, and the influence of motor control systems also need to be considered. If these factors are not strictly controlled, contradictory results may be obtained. Although there should be more than one read-out core area in the brain, there are far fewer of them relative to the various timing units distributed throughout the brain, and many modalities may even be sharing the same read-out system.

Multiple studies have reported that visual motion leads to perceptual time dilation[1, 42–45]. In contrast, we found that motion stimuli lead to less perceptual times. One reason for the different results is that these studies used a direct comparison of a static stimulus and a moving stimulus without considering that the populations of neurons they activate are actually different. In our study, the durations and spatial locations of the visual stimuli presented were the same, implying that the retinal ganglion cells activated under different stimulus conditions are identical. The only difference was the relationship between their spatial location over time, making motion perception available in one case and not in the other. Another reason could be that most of these studies had the subjects reproduce the durations after presenting a stimulus and then did the comparisons by the experimenters. The reproduction procedure itself introduces other factors, such as differences in memory extraction and perceptually guided behavior. By adopting an approach where the comparison is completed within the same trial and then the results of the perceptual comparison are reported through behavior, we can largely avoid the influence of the above factors.

### The limitations of current work

First, it should be clarified that our perceptual assembly model is mainly used to explain the neural basis of longer or shorter perceptual time at fixed physical time, rather than describing the variation pattern of perceptual time with physical time. Second, the current work is based on sensory time perception of visual signals in milliseconds to seconds range, it maybe not applicable to time perception when no sensory signal is available or when cross-modal comparisons are performed. Third, it’s a feedforward model and we did not considering any cross-modality regulation signals like attention or emotion. Attention may play an important role in perceptual state transitions or selection, e.g., attention can change the composition of a time estimation system by selecting perceptual states. Unfortunately, the present study was not able to provide evidence for this hypothesis. Additionally, the effect of neuroplasticity on time perception is not given a specific explanation, although it influences both time units and read-out rules. The reason for this is that we wanted to give a relatively simple and stable model framework by excluding plasticity, so that the neural mechanisms of time estimation and its covariation with visual stimuli are clearer. From another perspective, the stability of the performance of the time estimation system is very important. Even if stimulus-induced short-term and local plasticity has an effect, it should be limited.

## Methods

### Materials

In all experiments, the experimental programs were controlled using a Dell Vostro 5090 desk computer. Visual stimuli were generated using MATLAB with NIMH MonkeyLogic [46] and presented on a 32-inch Display++ (resolution, 1920 × 1080 pixels; refresh rate, 120 Hz). Participants were seated 55 cm away from the display and rested their heads on a chinrest to maintain a stable head position.

### Stimulus

The visual stimulus was a white dot (radius = 0.3°, [255 255 255]) presented at different spatial locations along a circular path (radius = 3°) centered at the screen’s center. Each stimulus lasted 3000 ms with 360 frames. The visual stimuli primarily included two types: ordered and randomized stimuli. In the ordered stimulus, the dot was sequentially presented at adjacent positions (clockwise or counterclockwise) to create the motion perception (apparent motion sequence, see Fig. 1a). In the randomized stimulus, the dot was randomly presented at pre-generated spatial positions. In both types of stimuli, the dot first appeared at the bottom of the circle.

The potential positions of the dot were determined by the event frequency, which was the number of positions where the dot appeared per second. The positions were calculated using equations (1) – (3), where *Fre*, *x*_F,j_, *y*_F,j_, and *R* represented the event frequency, horizontal coordinate, vertical coordinate, and the radius of the circular path, respectively. The variable *C* indicated the direction of motion, with +1 indicating clockwise and -1 indicating counterclockwise.

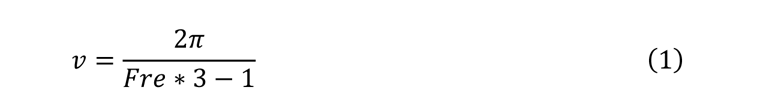

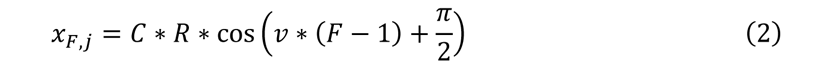

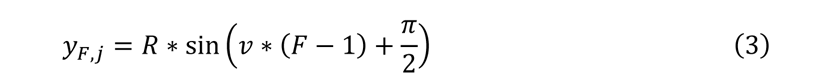

### Procedure

All participants were right-handed and had normal or corrected-to-normal visual acuity. Before the experiment, they all gave informed consent in accordance with procedures and protocols approved by the institutional review board of the Institute of Psychology, Chinese Academy of Sciences. All participants were naïve to the purpose of the experiments and received monetary compensation after the experiment.

The study comprised three experimental tasks. The first task, the duration comparison task, involved comparing the durations of two sequentially presented visual stimuli, with a standard duration of 3000 ms (see Fig. 1b). Each trial began with a red fixation cross (0.3° × 0.3°, [255 0 0]) for 300 ms, which changed to a white fixation point (0.3° × 0.3°, [255 255 255]) when the stimulus appeared. Participants were instructed to fixate on the fixation point throughout the experiment. Prior to the presentation of each stimulus, 300 ms of noise and 200 ms of a black screen with a fixation cross were displayed to ensure that the visual system was in a consistent state. The noise was created by randomly selecting 18 images from a set of 100 noise images, which were generated using the *MaskeTexture* and *DrawTexture* functions of Psychtoolbox [47, 48] in MATLAB (The MathWorks Inc., Natick, MA, USA). Each image filled the entire screen and was presented for two frames, resulting in dynamic random noise lasting 300 ms. After both visual stimuli were presented, participants were required to report which interval (the first or second) appeared longer within 3000 ms by pressing the left or right arrow keys, respectively. If no response was made, the trial would be repeated later. The intertrial interval (ITI) was 2000 ms. Before the experiment, participants were informed that there would be a slight temporal difference between the two stimuli. Additionally, to test whether participants were attentive during the task, we included catch trials with a genuine temporal difference (3000 ms vs. 2500 ms), accounting for 25% of all trials. Participants with an accuracy below 60% on these trials were excluded from the analysis due to lack of engagement.

The continuous motion discrimination task was conducted in Exp. 5a and 5b (see Fig. 4a and 4b). In each trial, we presented an ordered stimulus (as shown in Fig. 1a) with different event densities, which could be 15, 20, 24, 30, 40, 60, and 120 Hz. Before the stimulus presentation, 300 ms of noise and 200 ms of a black screen with a fixation cross were presented, similar to the duration comparison task. Participants were instructed to report whether they perceived the visual stimuli as continuous motion or flicker within 3 seconds by pressing the left or right arrow keys, respectively. In fact, as the event frequency increased, the spacing between the dot positions decreased, making the visual stimulus more likely to be perceived as continuous motion rather than flicker.

The difference threshold measurement task was conducted in Exp. 5b (see Fig. 4c). The visual stimuli used in this task were identical to those used in the duration comparison task. However, participants were instructed to judge whether there was any difference between two sequentially presented stimuli, regardless of whether the difference was related to time, motion, or any other factor. In addition, we included catch trials that involved a genuine temporal difference, which accounted for 25% of all trials. Participants whose accuracy was below 60% on these trials were excluded.

### Data Analysis

In the duration comparison task, we computed the probability of choice (POC) for each condition, defined as the probability that the condition was selected as longer in the trials that included the condition. We then performed one-way repeated-measured ANOVA on the POCs, and adopted Greenhouse-Geisser correction if the Mauchly’s sphericity assumption was violated. The p-values of *post-hoc* tests were Holm-corrected.

In the continuous motion discrimination task, we computed the identification rate (IR) for each condition, defined as the probability that the condition was perceived as continuous motion in the trials where it was presented. For each participant, we fitted the IR with a cumulative gaussian distribution (see equation (4)) and calculated the coefficient of determination (R-squared) and standard error (SE). The sensory threshold for continuous motion perception was set at the point where IR equaled 0.75. If the IR of a condition was greater than 0.75, the condition was perceived as continuous motion rather than flicker (see Fig. 4a and 4b).

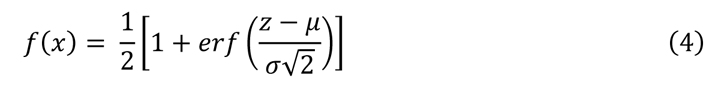

In the difference threshold measurement task, we computed the discrimination probability, which represents the probability of correctly distinguishing the difference between two sequentially presented stimuli. We performed related samples Friedman tests and pairwise comparisons to determine whether there were differences in discrimination ability among different conditions. Additionally, we performed one-sample Wilcoxon signed-rank tests on the median discrimination probability for each condition to determine whether there were significant differences between the discrimination probability and 0.25. A discrimination probability significantly higher than 0.25 indicated that the participants could effectively distinguish between the two stimuli, while a probability of 0.25 or lower suggested that they were making random choices.

All data analysis was performed using MATLAB 2017b and IBM SPSS statistics 27.

**Exp.1 (**Fig. 1c): Twenty-two participants (11 females; aged 23.23 ± 3.09 years, mean ± SD) joined the experiment. They completed a duration comparison task with ordered stimuli, which involved four different stimulus conditions (as shown in Fig. 1c): (1) the spatial location condition (20 Hz), where the dot repeatedly appeared twice at each position of 60 positions over three seconds; (2) the energy density condition (60 Hz), where the repetition times increased to six times at each position; (3) the color condition (60 Hz), similar to the repetition condition, but the dot’s color changed every two frames in the order blue/green/red; (4) the expanded spatial location condition (60 Hz), the number of positions increased to 180. In each trial, the conditions of the two ordered stimuli were different and randomly selected from the four conditions, resulting in six condition combinations (1-2, 1-3, 1-4, 2-3, 2-4, 3-4). The presentation order and the moving direction were balanced across trials. Therefore, each participant needed to complete 320 trials (6×2×2×10÷(1-0.25)=320). We computed each condition’s probability of choice (POC) and then performed one-way repeated-measured ANOVA on the POCs.

**Exp.2 (**Fig. 2a): Thirty participants (13 females; aged 23.50 ± 2.45 years) joined the experiment. Participants completed a duration comparison task with randomized stimuli, which involved three different stimulus conditions: 30 Hz (four repetitions of dots at 90 positions), 40 Hz (three repetitions of dots at 120 positions), and 60 Hz (two repetitions of dots at 180 positions). All types of visual stimuli were similar to those used in Exp.1, where the dot’s potential spatial locations were uniformly distributed. However, in the present experiment, the dot was randomly presented at all potential spatial locations, rather than being sequentially presented at adjacent spatial locations. In each trial, the event densities of the two stimuli were different and randomly selected from the three conditions. The presentation order of the two event densities was counterbalanced across trials, resulting in 80 trials (3×2×10÷(1-0.25)=80) in this experiment. We conducted a one-way repeated-measured ANOVA on the POC for each event frequency.

**Exp.3 (**Fig. 3a): Participants were the same in Exp. 2. Participants completed a duration comparison task. Each trial involved two visual stimuli: one ordered stimulus and one randomized stimulus, with the same event frequency, which could be 30 Hz, 40 Hz, and 60 Hz. Although the potential spatial locations of both types of visual stimuli were the same, the ordered stimulus involved spatiotemporal integration, while the randomized stimulus did not. The presentation order of the two stimuli types was counterbalanced across trials, resulting in 80 trials (3×2×10÷(1-0.25)=80) in this experiment.

We conducted a one-way repeated-measured ANOVA on the POCs of each event frequency, and found no difference across three conditions. Next, we compared the POCs of the randomized conditions with 0.5 under three event densities, using the one-sample t-test with Welch’s correction.

**Exp.4a (**Fig. 4a): Thirty-four participants (16 females; aged 23.24 ± 2.78 years) joined the experiment. Participants completed a duration comparison task and a continuous motion perception task. In the duration comparison task, visual stimuli consisted of four ordered stimuli with event densities of 15 Hz, 30 Hz, 60 Hz, and 120 Hz. The presentation order and the moving direction were balanced across trials. Therefore, each participant needed to complete 320 trials (6 x 2 x 2 x 10 + (1 -0.25) = 320). We computed the POC for each condition and conducted a one-way repeated-measured ANOVA on the POCs. We also computed the sensory threshold for continuous motion perception.

**Exp.4b (**Fig. 4b): Experimental procedures were same as Exp.4a. Twenty-two participants (11 females; aged 23.18 ± 2.32 years) joined the experiment. In the duration comparison task, visual stimuli consisted of three ordered stimuli with event densities of 30 Hz, 40 Hz, and 60 Hz. The presentation order and the moving direction were balanced across trials, resulting in 12 experimental conditions. Data analysis was the same as in Exp. 4a.

**Exp.4c (**Fig. 4c): Visual stimuli were the same as in Exp. 4b, but the behavioral task was different. It measures the non-temporal perceptual difference. Thirty-four participants were involved. Seven of them were excluded because of their accuracy below 60% in the deception trials, leaving 27 participants (15 females) for analysis (aged 22.81 ± 2.83 years). This task included three conditions: 30 Hz vs. 40 Hz, 40 Hz vs. 60 Hz, and 60 Hz vs. 120 Hz. Participants were required to determine whether the two sequentially presented stimuli were identical. The moving direction and the presentation order were balanced across trials, resulting in 12 experimental conditions. We conducted related samples Friedman tests and one-sample Wilcoxon signed-rank tests to investigate whether there were differences in discrimination ability among different conditions and whether participants could correctly distinguish between two stimuli.

**Exp.5a (**Fig. 6d): Twenty-two participants joined the experiment (11 females; aged 23.23 ± 3.09 years). The participants were required to perform a duration comparison task, which included three types of visual stimuli: constant speed motion, uniformly accelerating motion with an initial speed of zero, and uniformly decelerating motion with an initial speed of zero. The constant speed motion stimulus was the same in Exp. 1. For the uniformly accelerating motion stimulus, the dot’s positions were calculated by equations (5) – (7).

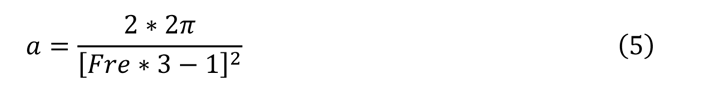

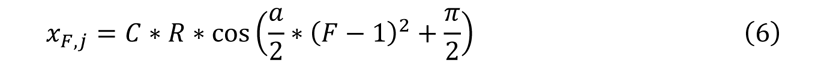

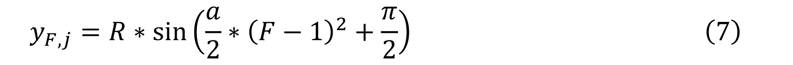

For the uniformly decelerating motion stimulus, the dot’s positions were calculated by equations (8) – (11).

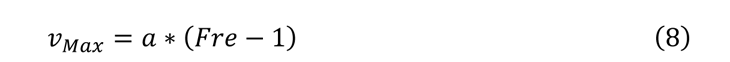

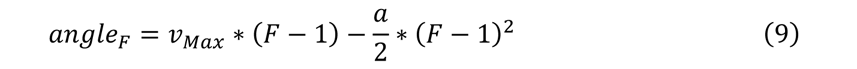

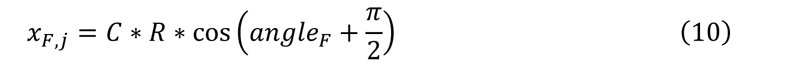

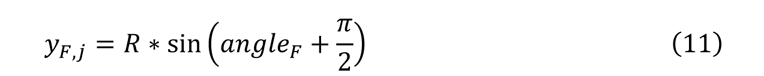

The event frequency of all three stimuli was set to 30 Hz. In each trial, two types of stimuli were randomly selected from them, resulting in three possible combinations (acc. vs. dec., acc. vs. con., dec. vs. con.). The presentation order and the moving direction were balanced across trials, resulting in 160 trials (3×2×2×10÷(1-0.25)=160) in the experiment. We performed a one-way ANOVA with *post-hoc* t-test on the POCs of each type of stimulus.

**Exp.5b&c (**Fig. 6e): This experiment differed from Exp.5a in the types of visual stimuli used. In Exp.5a, the visual stimuli consisted of uniformly accelerating or decelerating motion stimuli, which resulted in some spatial locations having smaller intervals due to the uniformity of the speed changes. This could have caused some of the spatial locations not to be perceived as a new independent event, leading to fewer events being perceived in the accelerating or decelerating motion stimuli than the constant speed motion stimulus. Therefore, in this experiment, we divided the circular path into several sections, with the spatial intervals being the same within each section but different between different sections. This experiment involved three types of visual stimuli: accelerating motion, decelerating motion, and constant speed motion. In the accelerating motion stimulus, the spatial interval of each section increased from small to large. Conversely, in the decelerating motion stimulus, the spatial interval of each section was in reverse order. In the constant speed motion stimulus, the number of positions was the same as in the accelerating and decelerating motion stimuli.

This experiment consisted of two conditions: a high spatial frequency condition and a low spatial frequency condition. In the high spatial frequency condition, twenty-nine participants (15 females; aged 25.17 ± 3.02 years) were recruited. The circular path (i.e., 360°) was divided into seven sections, with each section having a different radian and event frequency: 28°and 30Hz, 35°and 24Hz, 42° and 20Hz, 56° and 15Hz, 60° and 12Hz, 60°and 10Hz, and 75° and 8Hz. There was a total of 45 spatial positions. In the low spatial frequency condition, twenty-six participants (14 females; aged 24.92 ± 2.77 years) were recruited. The circular path was divided into five sections, with each section having a different radian and event frequency: 90° and 8Hz, 60°and 10Hz, 70° and 12Hz, 56°and 15Hz, 84°and 20 Hz. There were 40 spatial locations in total. The presentation order and moving direction were balanced across trials. The data analysis was the same as described before.

## Acknowledgements

We thank Prof. Yan Yang for valuable suggestions.

## Author contributions

T.Z. and Y.J. designed the study. W.H. developed the experimental paradigms, collected most of the data and did all the analysis. T.F. and Y.S. collected partial data. W.H. and T.Z. interpreted the results and prepared the figures. T.Z., W.H. and Y.H wrote the manuscript.

## Data Availability

Data available on request from the authors.

